# Longitudinal in vivo MR elastography reveals whole-liver viscoelastic involvement in a murine model of hepatocellular carcinoma

**DOI:** 10.1101/2025.09.04.674138

**Authors:** Pedro Augusto Dantas de Moraes, Yasmine Safraou, Karolina Krehl, Akvile Häckel, Tobias Haase, Susanne Metzkow, Rafaela Vieira da Silva, Yalei Zang, Jonas Röder, Anja A. Kühl, Lynn Jeanette Savic, Jürgen Braun, Eyk Schellenberger, Ingolf Sack, Jing Guo

## Abstract

Cancer cells actively shape their microenvironment and adapt the physical and biomechanical properties of the host tissue. However, for organs like the liver, especially in vivo, it remains unclear at what rate and spatial extent macroscopic viscoelastic properties change during formation of a cancer-permissive environment. Using clinical multifrequency magnetic resonance elastography (MRE) in a mouse model of hepatocellular carcinoma (HCC), we identified surprisingly early and large-scale viscoelasticity changes leading to whole-liver biomechanical involvement stretching far beyond the local cancer microenvironment. Widespread liver softening began two weeks after HCC inoculation, followed by a decrease in tissue viscosity and fluidity two weeks later, preceding any macroscopic evidence of local tumor growth. In contrast, local lesions with stiff-rigid biomechanical properties were not detectable by standard MRI and MRE until five to six weeks post-injection. Furthermore, tumor viscoelasticity correlated with that of the host liver, also suggesting a possible widespread adaptation of the biomechanical properties beyond the tumor margins and its local niche during early liver colonization. The observed large-scale viscoelastic signature, detectable just two weeks after tumor cells injection, could serve as a non-invasive imaging biomarker to inform physicians about tumor niche formation and liver cancer progression long before any macroscopic manifestation of solid tumors.

## Introduction

Liver cancer continues to present a significant global health issue, ranking as the sixth most prevalent cancer worldwide and the third leading cause of cancer-related mortality, mostly related to hepatocellular carcinoma (HCC), 75%–85% of cases^1^. This high mortality rate is attributable to a confluence of factors, including late-stage diagnosis, the aggressive nature of the disease, and the limited efficacy of current treatment options in advanced stages^2^.

Major risk factors for HCC include chronic alcohol consumption, metabolic dysfunction-associated liver disease, and viral infection^3^. Independent of the underlying etiology, neoplastic lesions typically develop a pre-stressed environment of chronic necroinflammation, which progresses from fibrosis to cirrhosis and, ultimately, HCC^4^. This cascade, originating from alterations in the extracellular matrix (ECM), indicates the critical role of the host tissue in the proliferation of cancer cells and the formation of a cancer-permissive environment.

One fundamental guiding principle of tumor progression is the exchange of mechanosensory signals between cells and ECM^5,6^. This biomechanical crosstalk allows cancer cells to actively shape their microenvironment and adapt to the physical and biomechanical properties of their host tissue^7^. In addition, the collective behavior of cancer cells influences the coarse-grained material properties of solid tumors and provides a macroscopic link to the emergent state of cancer cells in correlation with their aggressiveness and invasive behavior^8,9^. Therefore, measurement of in vivo biomechanical properties of HCC and host tissues by elastography techniques not only provides a new imaging contrast complementary to established multiparametric magnetic resonance imaging (mp-MRI), but also opens a window into the microscopic biomechanical structures that change during the development, progression and invasion of liver tumors^10^. In current clinical routine, multiparametric MRI enables early detection of small HCC lesions in patients with cirrhosis or fibrosis^11^. However, its prognostic value remains limited in identifying precancerous niches or the development of a tumor-permissive microenvironment before macroscopic tumors become visible¹¹. While magnetic resonance elastography (MRE)^12^ is still in its early stages with respect to high-resolution spatial mapping of stiffness, attenuation, and tissue fluidity^13^, current reports highlight the potential value of MRE as a clinical imaging modality in liver tumors^14^, specifically for the differentiation of tumor malignancies^15^, the detection of HCCs with glypican-3 (GPC3) expression^16^, the investigation of microvascular invasion^17–20^ pathological classification,^21–23^, and prognosis^24–28^.

As a tomographic modality, MRE covers both tumor and tumor-adjacent tissue which facilitates studies of tumor-niche biomechanical interactions in vivo^29^. In HCC, a more infiltrative tumor behavior has been observed in stiffer compared to softer livers^30^. This observation highlights the need to delineate the relative contributions of precancerous liver states versus cancer cell–intrinsic activity to changes in liver mechanical properties and the establishment of a permissive mechanical niche that drives HCC proliferation and invasiveness. If HCC affects liver tissue on a larger scale beyond the tumor microenvironment, MRE may be able to detect subtle biomechanical hallmarks of tumor development in apparently unaffected tissue prior to the visible manifestation of solid tumors in mp-MRI. To test this hypothesis under controlled conditions in vivo, we implanted HCC into the livers of mice and analyzed biophysical tissue changes using longitudinal mp-MRI and novel multifrequency MRE. As the study protocol was developed on a clinical MRI system, the identified imaging markers can be easily translated into diagnostic MRE and mp-MRI.

## Results

Our mp-MRI and MRE imaging protocol provided T2-weighted images (T2w), as well as biophysical parameter maps of shear wave speed (SWS), penetration rate (PR), loss angle (*φ*) and apparent diffusion coefficient (ADC) as proxies for tissue stiffness, inverse viscosity (shear wave penetration), tissue fluidity and water diffusion, respectively, as further detailed in the Methods section. In vivo MR images were acquired for each time point: baseline, two weeks (2w), four weeks (4w) and five to six weeks (5/6w) post tumor inoculation.

### Tumor lesions are detectable by MRI only at late time points

Tumors were not clearly macroscopically discernible on T2w images in all 26 mice until 5/6w after implantation. The progression of the tumor in one mouse can be seen in Fig. 1, displaying representative T2w, MRE magnitude, SWS, PR, and ADC images for all four time points. Liver SWS declined from 2w onwards, while liver PR increased particularly at 5/6w when the tumor became visible. At 5/6w, the tumor exhibited higher SWS, PR, and ADC values than the liver. Alteration in fluidity was observed with a decrease in *φ* at the last time point, for both liver and tumor.

**Fig. 1:**
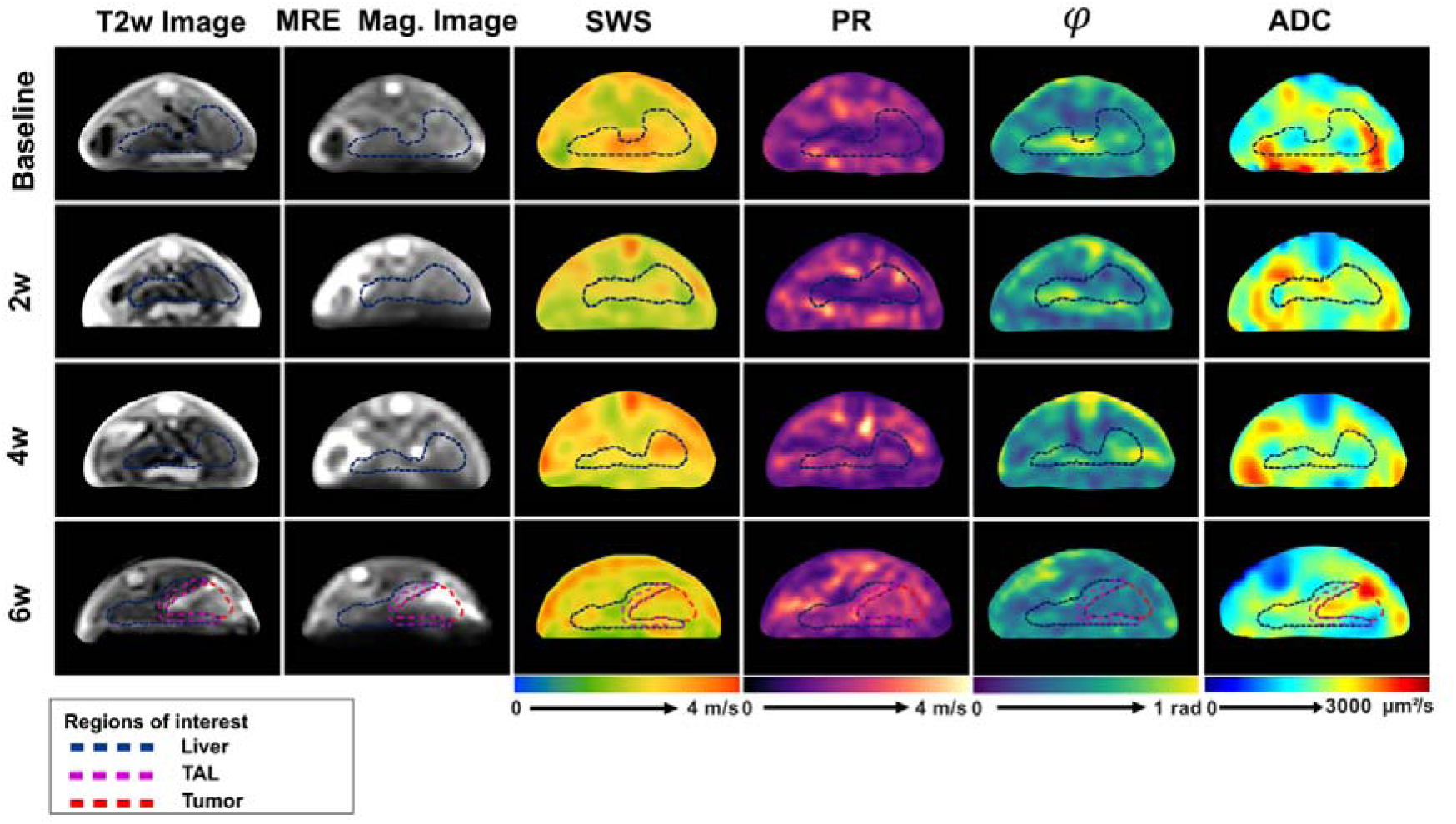
Representative longitudinal changes of MRI, MRE and ADC maps of the liver, tumor, and tumor-adjacent liver (TAL) regions at baseline, 2 weeks (2w), 4 weeks (4w), and 6 weeks (6w) post cancer cells inoculation. Columns display T2-weighted (T2w) anatomical images, MRE magnitude images, shear wave speed (SWS), penetration rate (PR), loss angle (cp), and apparent diffusion coefficient (ADC) maps. Regions of interest for the liver, TAL, and tumor are outlined by blue, purple, and red dashed lines, respectively. SWS maps provide stiffness measurements, while PR maps indicate inverse viscosity, φ maps reflect tissue fluidity, and ADC maps depict water diffusion. Imaging data reveal dynamic changes in tumor and liver biomechanics over time.

### Tumor progression shifts liver to soft-solid properties

Repeated measures ANOVA analysis of the imaging parameters (Fig. 2) revealed a reduction in liver SWS over the course of tumor development (*p <* 0.0001), already observed after two weeks (baseline: 2.7 ± 0.1 m/s vs. 2w: 2.6 ± 0.1 m/s, *p =* 0.02). Liver SWS reduced by 9.7 ± 0.6% at 5/6w compared to baseline values (baseline: 2.7 ± 0.1 m/s versus 5/6w: 2.4 ± 0.1 m/s, *p <* 0.0001) while liver PR increased by 17.7% ± 4.4% over the entire time course (baseline: 1.4 ± 0.2 m/s versus 5/6w: 1.7 ± 0.2 m/s, *p <* 0.0001). Pairwise comparison also revealed a significant increase in liver PR compared to baseline after 4w (1.6 ± 0.2 m/s versus 1.7 ± 0.2 m/s, *p =* 0.023). Liver ADC also reduced by 19.6% ± 6.9% (baseline: 1617 ± 215 µm²/s versus 5/6w: 1283 ± 329 µm²/s, *p <* 0.01). Additionally, at 5/6w, liver *φ* decreased by 19.9% ± 5.7% (baseline: 0.6 ± 0.1 rad vs. 5/6w: 0.5 ± 0.1 rad, *p <* 0.0001), suggesting an alteration in tissue fluidity during tumor progression.

**Fig. 2:**
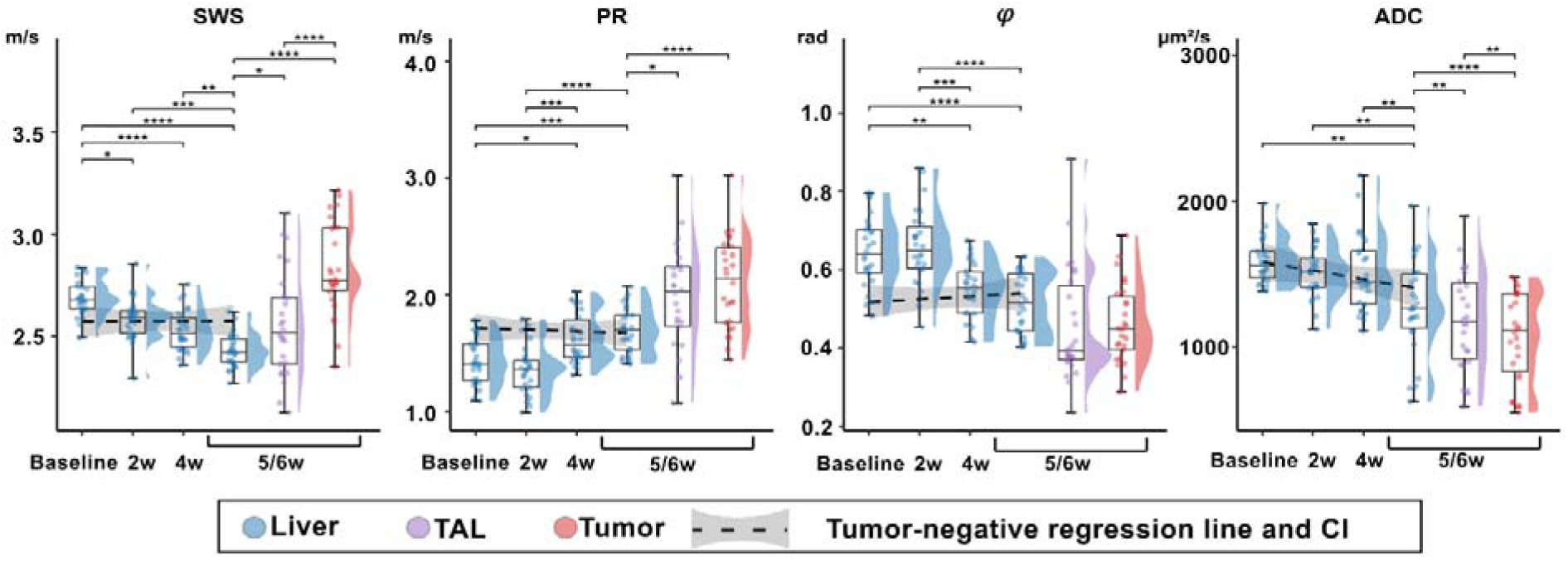
Longitudinal changes in imaging biomarkers across liver, tumor-adjacent liver, and tumor regions during tumor progression. Combined raincloud plots depicting longitudinal changes in shear wave speed (SWS), penetration rate (PR), phase angle (cp), and apparent diffusion coefficient (ADC) in the liver (blue), tumor-adjacent liver (TAL, magenta), and tumor (red) at different time points during tumor development. The tumor-negative group is represented by a regression line with a confidence interval (CI) shown in gray. Statistical analysis was performed using repeated-measures one-way ANOVA followed by post-hoc tests, with significance levels indicated as \*\*\*\**p<*0.0001, \*\*\**p<*0.001, \*\**p<*0.01, and \**p<*0.05.

### Tumors are stiffer and less attenuating than the host liver

Comparisons between regions of tumor, liver, and tumor-adjacent liver (TAL), were performed at 5/6w, when tumors were visible in the T2w images (Fig. 2). Tumor SWS and PR were higher than that of the liver (SWS: 2.8 ± 0.2 m/s versus 2.4 ± 0.1 m/s, *p <* 0.0001, PR: 2.0 ± 0.4 m/s versus 1.7 ± 0.2 m/s, *p <* 0.0001) while tumor ADC showed reduced values compared to liver ADC (1072 ± 306 µm²/s versus 1283 ± 326 µm²/s, *p <* 0.0001), revealing that tumors are characterized by stiffer and less viscous tissue properties with reduced water diffusivity than the host liver.

TAL exhibited intermediate biophysical properties, characterized by reduced SWS (2.5 ± 0.3 m/s, *p* < 0.0001) and comparable PR (2.0 ± 0.5 m/s, *p* = 0.7) compared to tumor tissue. Additionally, TAL demonstrated higher SWS (*p* = 0.05) and PR (*p* = 0.01) compared to the remaining liver parenchyma. Correspondingly, TAL ADC was higher than in tumors (1178 ± 341 µm²/s, *p* < 0.01) while lower than in the liver (*p =* 0.001). With *φ* = 0.5 ± 0.1 rad, tissue fluidity was not altered across tumor-liver-TAL regions, suggesting adaptations of whole-liver fluidity during HCC progression.

### Biophysical properties of the tumor-negative liver remain unchanged over time

At 5/6w, 10 mice which received HCC cells injection didn’t develop visible tumors, constituting a tumor-negative group. Hematoxylin and Eosin (H&E) stainings of their tissue samples showed micro scars and fibrotic regions without viable tumor cells. As demonstrated in Table 3 cell density, protein markers (MMP-2, MMP-9, CD31, Ki-67, and Vimentin), and the amount of glycosaminoglycans (GAGs) and collagen in these tumor-negative livers, as assessed by quantitative histology, showed no significant differences when compared to livers from the health control group (all *p >* 0.05).

Furthermore, biophysical properties SWS, PR, *φ*, and ADC of the livers in this tumor-negative group showed no significant changes over time (all *p >*0.05, regression lines in Fig. 2 At baseline, biomechanical properties differed between later tumor-positive livers and tumor-negative according to SWS (2.6 ± 0.1 m/s, –4.40% ± 0.30%, *p <* 0.01), PR (1.7 ± 0.2 m/s, +20.78% ± 5.46%, *p <* 0.01) and *φ* (0.5 ± 0.1 m/s, –21.53% ± 6.02%, *p <* 0.0001)

All results pertaining to the group analysis are compiled in Table 1.

**Table 1:**
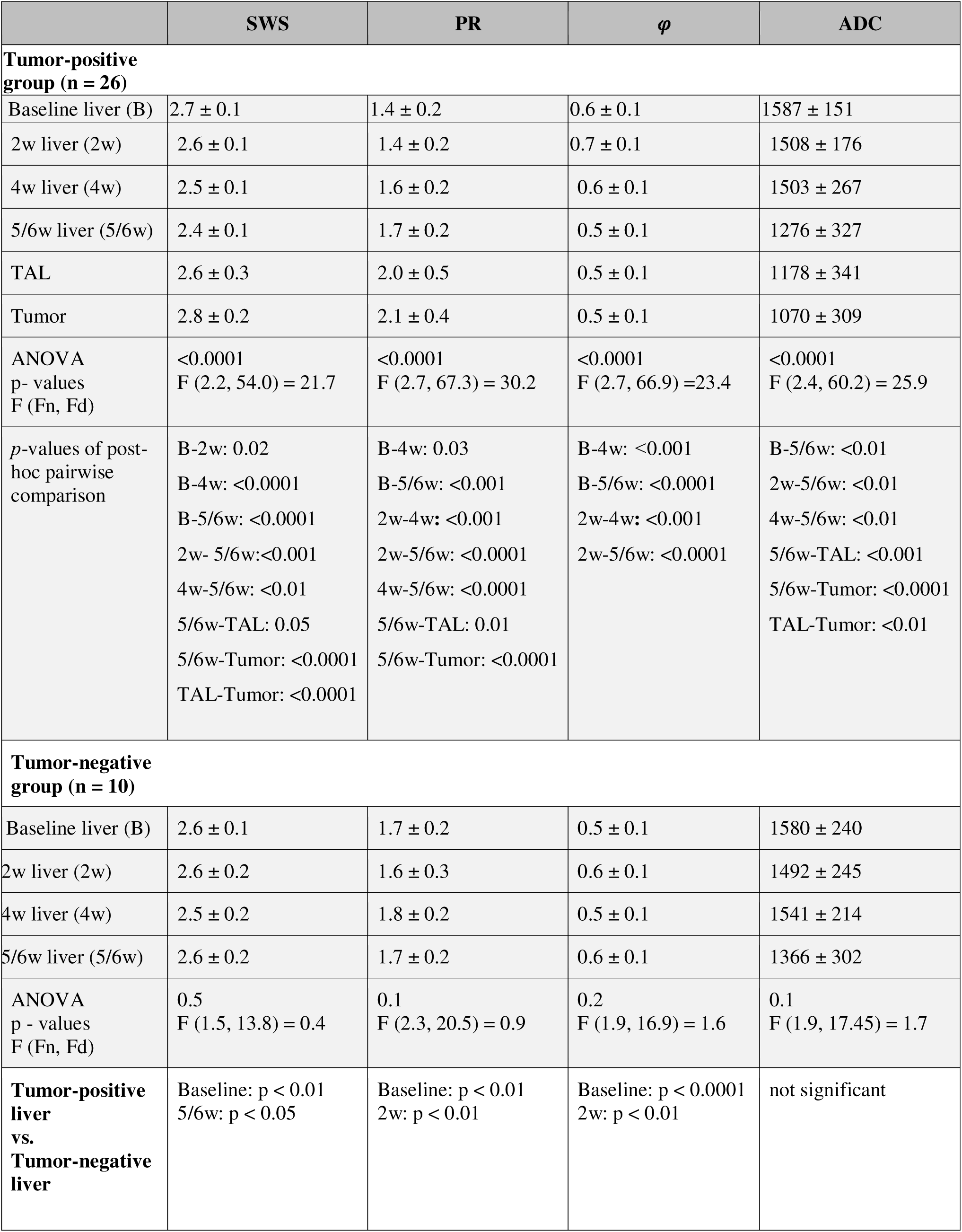
Group mean±SD values of imaging parameters shear wave speed (SWS), penetration rate (PR), and apparent diffusion coefficient (ADC) of liver, tumor, and tumor adjacent liver (TAL) were collected at different time points. For multiple comparison tests, p-values of ANOVA and the post-hoc pairs with significant difference were listed. F-statistics of the ANOVA test was reported as F (Fn, Fd), with Fn and Fd showing the degree of freedom of the numerator and the denominator, respectively. P-values comparing the tumor-positive and tumor-negative livers at different time points were also summarized.

| | SWS | PR | $\varphi$ | ADC |
| --- | --- | --- | --- | --- |
| **Tumor-positive group (n = 26)** |  |  |  |  |
| Baseline liver (B) | 2.7 ± 0.1 | 1.4 ± 0.2 | 0.6 ± 0.1 | 1587 ± 151 |
| 2w liver (2w) | 2.6 ± 0.1 | 1.4 ± 0.2 | 0.7 ± 0.1 | 1508 ± 176 |
| 4w liver (4w) | 2.5 ± 0.1 | 1.6 ± 0.2 | 0.6 ± 0.1 | 1503 ± 267 |
| 5/6w liver (5/6w) | 2.4 ± 0.1 | 1.7 ± 0.2 | 0.5 ± 0.1 | 1276 ± 327 |
| TAL | 2.6 ± 0.3 | 2.0 ± 0.5 | 0.5 ± 0.1 | 1178 ± 341 |
| Tumor | 2.8 ± 0.2 | 2.1 ± 0.4 | 0.5 ± 0.1 | 1070 ± 309 |
| ANOVA<br>p- values<br>F (Fn, Fd) | <0.0001<br>F (2.2, 54.0) = 21.7 | <0.0001<br>F (2.7, 67.3) = 30.2 | <0.0001<br>F (2.7, 66.9) = 23.4 | <0.0001<br>F (2.4, 60.2) = 25.9 |
| p-values of post-hoc pairwise comparison | B-2w: 0.02<br>B-4w: <0.0001<br>B-5/6w: <0.0001<br>2w- 5/6w: <0.001<br>4w-5/6w: <0.01<br>5/6w-TAL: 0.05<br>5/6w-Tumor: <0.0001<br>TAL-Tumor: <0.0001 | B-4w: 0.03<br>B-5/6w: <0.001<br>2w-4w: <0.001<br>2w-5/6w: <0.0001<br>4w-5/6w: <0.0001<br>5/6w-TAL: 0.01<br>5/6w-Tumor: <0.0001 | B-4w: <0.001<br>B-5/6w: <0.0001<br>2w-4w: <0.001<br>2w-5/6w: <0.0001 | B-5/6w: <0.01<br>2w-5/6w: <0.01<br>4w-5/6w: <0.01<br>5/6w-TAL: <0.001<br>5/6w-Tumor: <0.0001<br>TAL-Tumor: <0.01 |
| **Tumor-negative group (n = 10)** |  |  |  |  |
| Baseline liver (B) | 2.6 ± 0.1 | 1.7 ± 0.2 | 0.5 ± 0.1 | 1580 ± 240 |
| 2w liver (2w) | 2.6 ± 0.2 | 1.6 ± 0.3 | 0.6 ± 0.1 | 1492 ± 245 |
| 4w liver (4w) | 2.5 ± 0.2 | 1.8 ± 0.2 | 0.5 ± 0.1 | 1541 ± 214 |
| 5/6w liver (5/6w) | 2.6 ± 0.2 | 1.7 ± 0.2 | 0.6 ± 0.1 | 1366 ± 302 |
| ANOVA<br>p - values<br>F (Fn, Fd) | 0.5<br>F (1.5, 13.8) = 0.4 | 0.1<br>F (2.3, 20.5) = 0.9 | 0.2<br>F (1.9, 16.9) = 1.6 | 0.1<br>F (1.9, 17.45) = 1.7 |
| **Tumor-positive liver vs. Tumor-negative liver** | Baseline: p < 0.01<br>5/6w: p < 0.05 | Baseline: p < 0.01<br>2w: p < 0.01 | Baseline: p < 0.0001<br>2w: p < 0.01 | not significant |

### Tumor fluidity correlates with stiffness despite lower attenuation

Within the tumor, PR was negatively correlated with SWS (r = –0.4, *p* = 0.04) and ADC (*r =* –0.5, *p=*0.02), while *φ* showed a positive correlation with SWS (*r =* 0.7, *p <* 0.001) and ADC (*r =* 0.4, *p =* 0.03). Pooling data from all time points in the tumor positive group, liver SWS was positively correlated with liver ADC (*r =* 0.4, *p <* 0.0001) and negatively correlated with liver PR (*r =* –0.5, *p <* 0.0001). Additionally, liver *φ* exhibited a strong positive correlation with liver SWS (*r =* 0.6, *p <* 0.0001) and a weak positive correlation with liver ADC (*r =* 0.2, *p =* 0.05).

At 5/6w, tumor-liver properties were positively correlated for PR (*r =* 0.4, *p =* 0.02), *φ* (*r =* 0.5, *p =* 0.02) and ADC (*r=*0.8, *p<*0.0001) while only a weak positive correlation was observed for SWS (*r =* 0.4, *p =* 0.05).

The TAL, SWS correlated strongly with tumor and liver SWS (both *r =* 0.7, *p <* 0.0001), while PR and *φ* inversely correlated with that of tumor and liver (*r =* –0.9, *p <* 0.0001). TAL ADC closely aligned with liver ADC (*r =* 1.0, *p <* 0.0001).

Because *φ* and PR are not entirely independent biomechanical properties, they were highly correlated for liver (*r =* –0.9; *p <* 0.0001) and tumor (*r =* –0.9; *p <* 0.0001). The results pertaining to the correlation analysis are summarized in Table 2 and illustrated in Fig. 3.

**Fig. 3:**
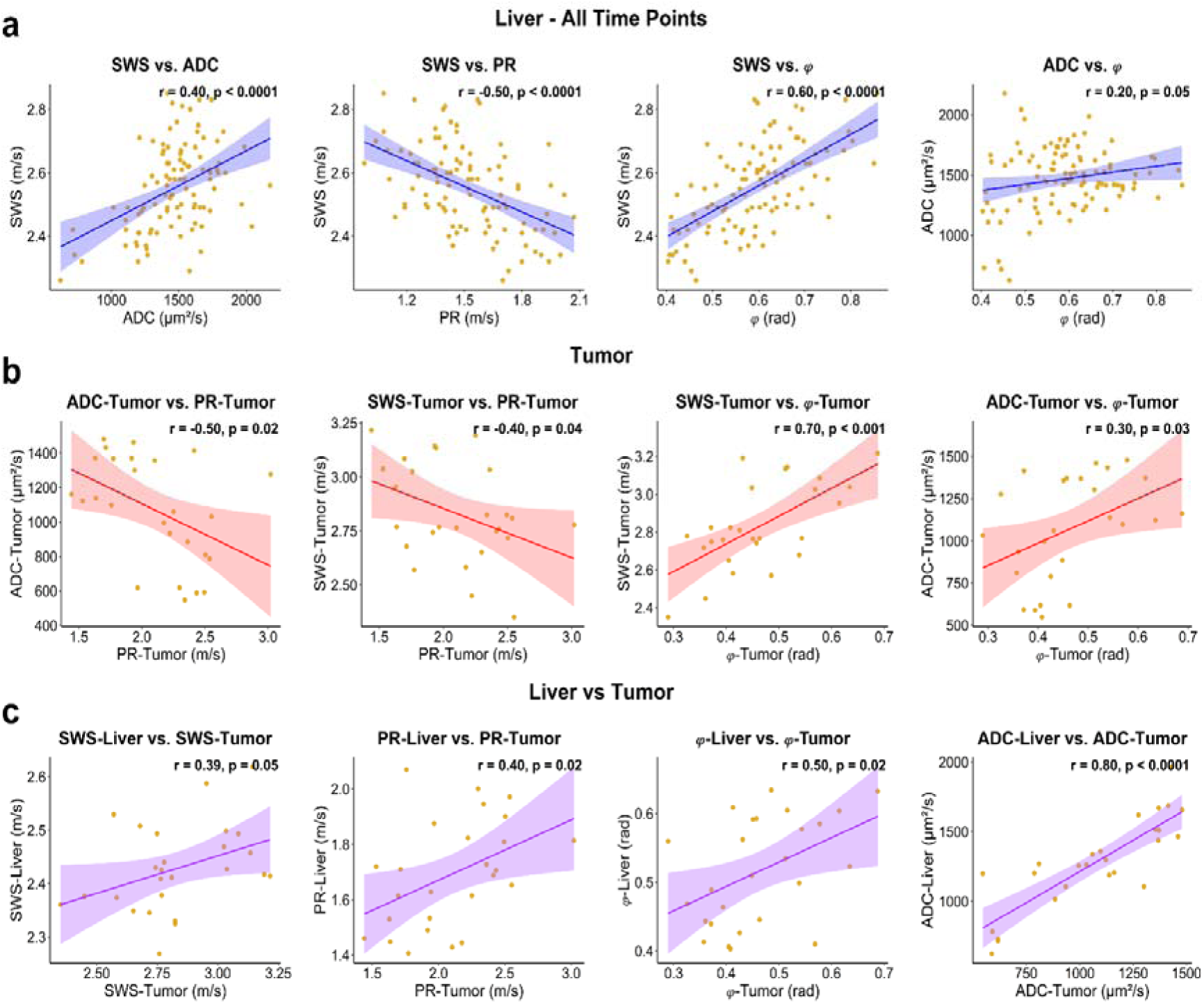
Correlation analysis of biomechanical parameters and water diffusion measured as shear wave speed (SWS), penetration rate (PR), phase angle (**<p**) and apparent diffusion coefficient (ADC). **a** Parameters of the full liver without tumor region pooling all values acquired at all time points from week 0 to week 6 after inoculating HCC cells. **b** Parameters of tumor tissue acquired from 5/6w when tumors became apparent. **c** Correlation analysis of parameters from the liver and tumor region at 5/6w. Each panel presents linear regression analysis with linear correlation coefficients (r) and p-values based on the Pearson correlation.

**Table 2:**
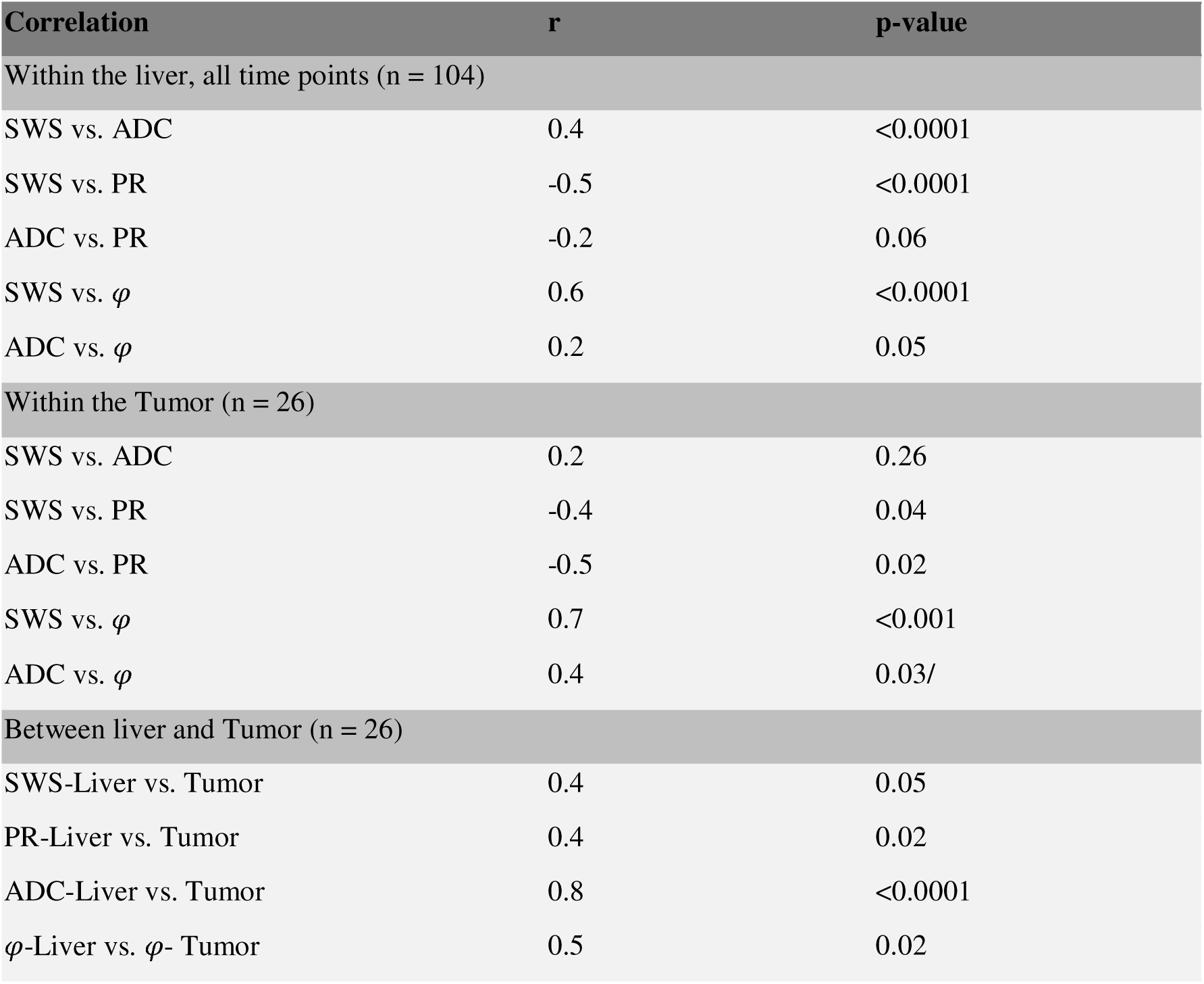
Results from the correlation analysis in the tumor-positive group between imaging parameters within the same region and between different regions for the same imaging parameters.

| Correlation | r | p-value |
| --- | --- | --- |
| Within the liver, all time points (n = 104) |  |  |
| SWS vs. ADC | 0.4 | <0.0001 |
| SWS vs. PR | -0.5 | <0.0001 |
| ADC vs. PR | -0.2 | 0.06 |
| SWS vs. $\varphi$ | 0.6 | <0.0001 |
| ADC vs. $\varphi$ | 0.2 | 0.05 |
| Within the Tumor (n = 26) |  |  |
| SWS vs. ADC | 0.2 | 0.26 |
| SWS vs. PR | -0.4 | 0.04 |
| ADC vs. PR | -0.5 | 0.02 |
| SWS vs. $\varphi$ | 0.7 | <0.001 |
| ADC vs. $\varphi$ | 0.4 | 0.03/ |
| Between liver and Tumor (n = 26) |  |  |
| SWS-Liver vs. Tumor | 0.4 | 0.05 |
| PR-Liver vs. Tumor | 0.4 | 0.02 |
| ADC-Liver vs. Tumor | 0.8 | <0.0001 |
| φ-Liver vs. φ- Tumor | 0.5 | 0.02 |

**Table 3:** Quantitative assessment of all histological markers as well collagen fibers characterizations across five anatomical regions: healthy liver, tumor-negative liver, tumor-positive liver, tumor-adjacent liver (TAL), and tumor. Multiple comparison tests were performed in the liver group (healthy liver, tumor-negative liver, and tumor-positive liver) and in the tumor-positive group (liver, TAL, and tumor), and the corresponding p-values were provided. Group statistics are presented as mean ± standard deviation or geometric mean (×/÷ geometric SD).

| Markers (units) | Healthy liver | Tumor-negative liver | Tumor-positive liver | TAL | Tumor | P-value of multiple comparison tests |
| --- | --- | --- | --- | --- | --- | --- |
| Nuclei density (counts/mm<sup>2</sup>) | 3785±550 | 3766±442 | 4539±1473 | 5106±1580 | 9301±3339 | Liver group:0.01<br>Tumor-positive group: <0.0001 |
| MMP-2 (counts/mm<sup>2</sup>) | 7.3(×/÷3.9) | 2.2(×/÷2.1) | 58.0(×/÷4.5) | 64.8(×/÷4.0) | 95.8(×/÷3.4) | Liver group:<0.0001<br>Tumor-positive:0.08 |
| MMP-9 (counts/mm<sup>2</sup>) | 7.57(×/÷3.6) | 32.5(×/÷1.6) | 54.48(×/÷3.4) | 62.60(±3.2) | 162.0(×/÷2.3) | Liver group:<0.001<br>Tumor-positive group:<0.0001 |
| CD-31 counts/mm<sup>2</sup> | 9.6 (×/÷ 5.0) | 8.6(×/÷1.5) | 30.14(×/÷2.0) | 46.29(×/÷1.7) | 99.7(×/÷1.9) | Liver group:<0.001<br>Tumor-positive group : <0.0001 |
| Ki-67 counts/mm<sup>2</sup> | 8.73(×/÷1.9) | 3.9(×/÷2.7) | 35.32(×/÷2.2) | 58.8(×/÷3.3) | 568.4(×/÷3.7) | Liver group: <0.0001<br>Tumor-positive group: <0.0001 |
| Vimentin (%) | 4.5(×/÷1.7) | 5.6(×/÷1.8) | 11.9(×/÷2.8) | 15.4(×/÷2.7) | 44.8(×/÷2.0) | Liver group: 0.007<br>Tumor-positive group :<0.0001 |
| Alcian blue (%) | 7.35(×/÷2.1) | 4.5(×/÷4.2) | 19.3(2.2) | 23.3(×/÷1.9) | 44.9(×/÷1.5) | Liver group:<0.001<br>Tumor-positive group:<0.0001 |
| Sirius Red (%) | 1.7(×/÷1.4) | 1.8(×/÷1.4) | 2.4(×/÷2.4) | 4.3(×/÷2.8) | 15.7(×/÷2.4) | Liver groups:0.41<br>Tumor-positive group:<0.0001 |
| Collagen coherency | 0.2±0.1 | 0.2±0.1 | 0.09±0.04 | 0.2±0.1 | 0.2±0.1 | Liver groups:0.0001<br>Tumor-positive group: <0.001 |
| Collagen red birefringent component (%) | 9.7±2.1 | 10.4±3.7 | 15.7±4.5 | 20.0±4.0 | 30.2±8.5 | Liver groups: <0.0001<br>Tumor-positive group: <0.0001 |
| Collagen yellow birefringent component (%) | 4.2±0.8 | 3.8±1.4 | 6.6±1.6 | 9.68±3.1 | 13.11±3.8 | Liver groups: <0.0001<br>Tumor-positive group:<0.0001 |
| Collagen green birefringent component (%) | 86.1±2.8 | 85.8±5.0 | 77.7±5.9 | 70.4±6.4 | 56.7±11.4 | Liver groups:<0.0001<br>Tumor-positive group:<0.0001 |

### Histopathology detects tumor-associated matrix remodeling in the liver

As shown in Fig. 4, H&E staining at 5/6 weeks post-inoculation revealed a well-demarcated separation between tumor and liver tissue, characterized by high nuclei density (counts/mm^2^) within the tumor (tumor: 9301 ± 3339 vs TAL: 5106 ± 1580 vs liver: 4539 ± 1473, *p* < 0.0001). Tumors showed a complex structure exhibiting a defined capsule, internal septae, and few necrotic zones. Alcian blue staining showed the presence of GAGs and acidic mucopolysaccharides in both tumor and host liver tissue, with higher staining-positive area (area percentage %) in the tumor than in the liver (tumor: 44.9 ×/÷ 1.5 vs liver: 19.3 ×/÷ 2.2, *p* < 0.0001. In addition, the GAG content was higher in tumor-positive livers than in control healthy livers (19.3 ×/÷ 2.2 % vs. 7.5 ×/÷ 2.3 %, *p* = 0.01). Sirius red staining demonstrated collagen deposition predominantly in the tumor capsule (15.7 ×/÷L2.4 %), while significantly lower amount of collagen was observed in distal liver parenchyma (2.4 ×/÷L2.4 %, *p* < 0.0001). Birefringence analysis of the collagen fibers revealed high mature collagen content (expressed in % of the total birefringent signal) in tumor (red: 30.19 ± 8.50% and yellow 13.1 ± 3.8%) than that of the surrounding liver (red: 15.7 ± 4.4% and yellow 6.60 ± 1.63%, *p* < 0.0001). Collagen fiber coherency obtained from gradient-based orientation mapping also showed that the fibers were more aligned in the tumor (0.17 ± 0.10) than in TAL (0.16 ± 0.10, *p* < 0.001) and the liver (0.9 ± 0.04, *p* < 0.001).

**Fig. 4:**
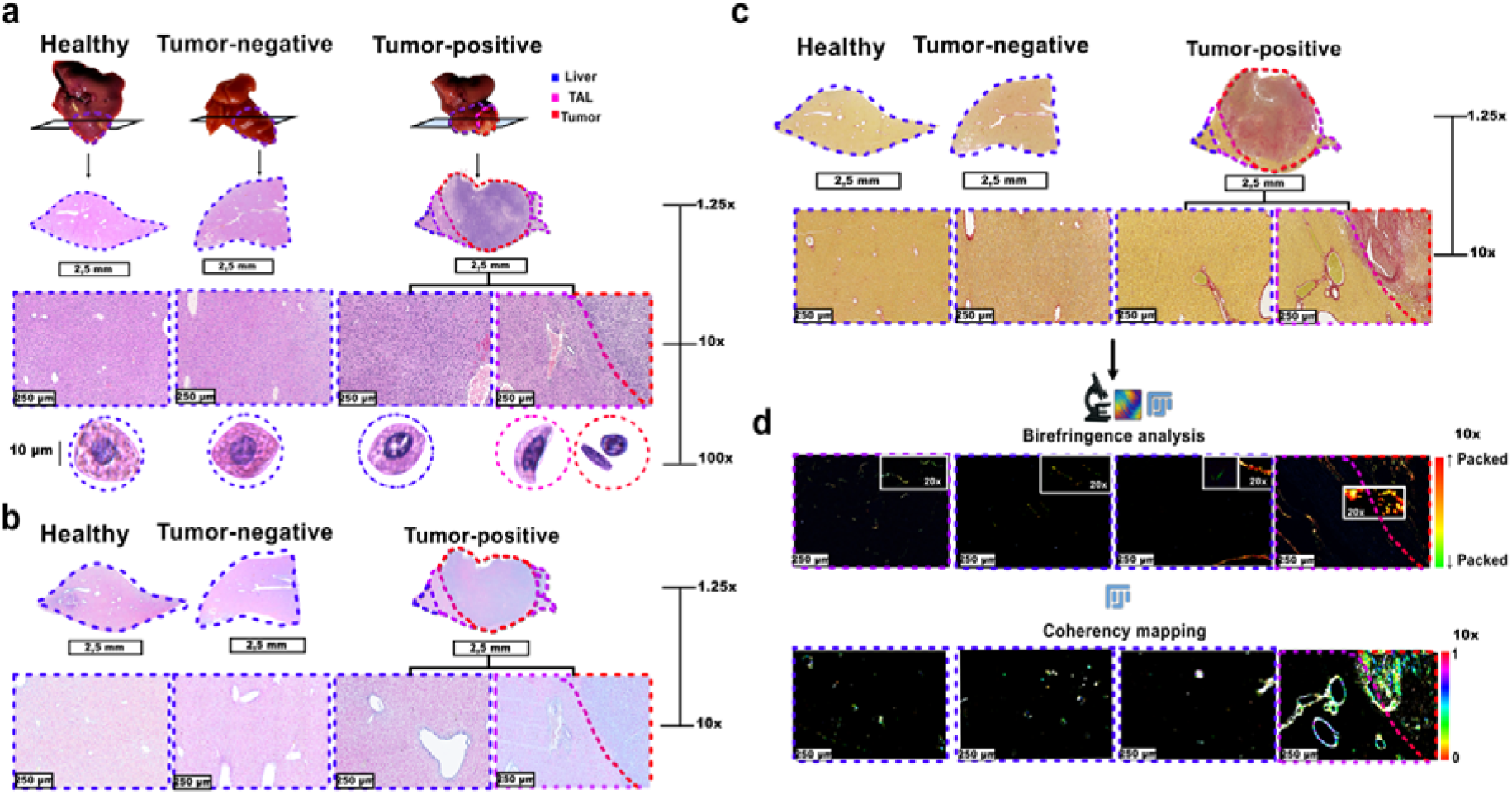
Multiscale histopathological and structural analysis reveals tumor-driven microarchitectural disruption and extracellular matrix remodeling in the liver. **Macroscopic** liver images demonstrate the histological slice orientation in healthy (left), tumor-negative (middle), and tumor-positive (right) animals. Representative hematoxylin and eosin (H&E) staining is shown at 1.25×, 10×, and 100× magnifications, highlighting liver (blue), tumor-adjacent liver (TAL, purple), and tumor (red) regions. H&E staining showed characteristic features of the tumor such as high nuclei density, irregular shape, and hyperchromatic nuclei. **b** Alcian blue staining reveals progressive accumulation of acidic polysaccharides and glycosaminoglycans, particularly at the tumor-liver interface. **c** Sirius red staining revealed dense collagen bundles enrichment in the tumor and tumor-adjacent areas. **d** Polarized light microscopy analysis of the same Sirius red-stained sections as shown in c. The presence of a red-dominated birefringent pattern, together with elevated levels of fiber coherency within the tumor region, signaled the deposition of dense, highly aligned collagen fibers. Insets show zoomed (20×) views.

In immunofluorescence images (Fig. 5), Ki-67 positive counts (counts/mm²) were elevated in tumor (568.4 ×/÷L3.7) which was significantly higher than in TAL (58.8×/÷L3.3) and tumor-positive liver (35.3 ×/÷L2.2, *p <* 0.0001). Similarly, CD31 expression (counts/mm²) was significantly higher in tumor areas (568.4 ×/÷L 3.7) compared to both TAL (58.8 ×/÷ 3.3) and tumor-positive liver (35.3 ×/÷L2.2, *p <* 0.0001). Vimentin was detected predominantly in tumor regions (44.8 ×/÷L2.0, area percentage %), including both intracellular and extracellular compartments, and to a lesser degree in surrounding liver tissue (11.9 ×/÷ L2.8, *p <* 0.01).

**Fig. 5:**
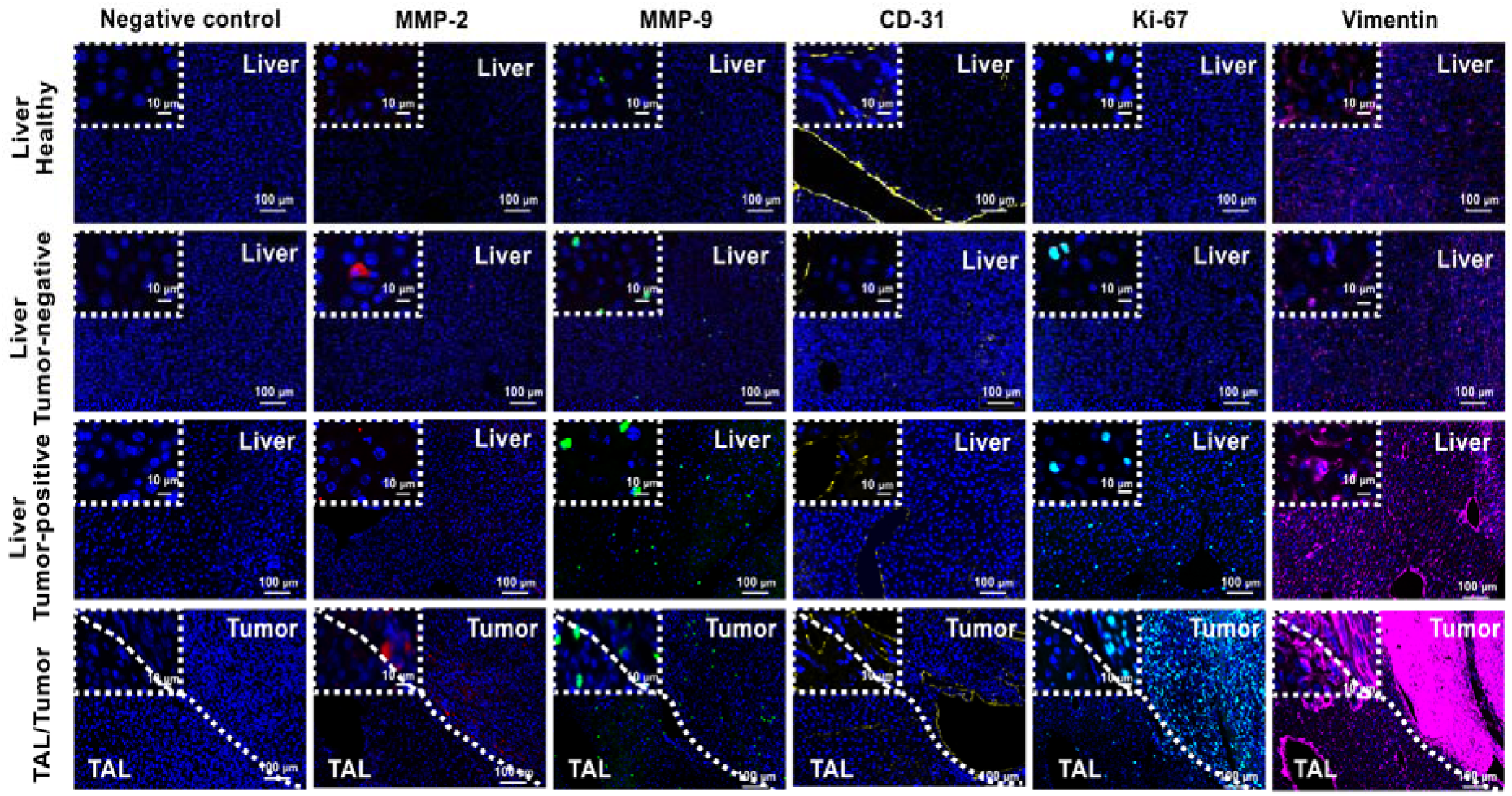
Immunofluorescence of tumor-liver transition zones reveals extracellular matrix remodeling, tumor cell infiltration, and angiogenesis in the tumor and surrounding tissue. Immunofluorescence staining for MMP-2 (red), MMP-9 (green), CD-31 (yellow), Ki-67 (cyan), and vimentin (magenta) in healthy liver, tumor-negative liver, tumor-positive liver, tumor-adjacent liver (TAL), and tumor. Images are shown at 10× magnification with 100× insets. Negative control staining was performed with only the secondary antibody to exclude nonspecific binding. Tumor and liver areas are separated by a white dotted line. The analysis highlights matrix remodeling, indicated by MMP-2 and MMP-9, increased proliferative activity (Ki-67), vascularization (CD-31), and mesenchymal features (vimentin) within the tumor microenvironment.

The presence of MMP-9 (counts/mm²) in the tumor (171.7 ×/÷ 2.0) was found to be significantly higher compared to that in the surrounding liver (54.5 ×/÷ 3.4) and the TAL (62.60 ×/÷ 3.20), *p* < 0.0001). Furthermore, the presence of both MMP-2 (58.0 ×/÷ 4.51, counts/mm²) and MMP-9 (54.5 ×/÷ 3.4, counts/mm²) identified in tumor-bearing livers exhibiting a marked increase in expression relative to the control healthy liver (MMP-2: 7.5 ×/÷ 3.9, *p* < 0.001 and MMP-9: 7.6 ×/÷3.6, *p* < 0.01, respectively). The results of the group analysis of the quantitative histological data are compiled in Table 3 and illustrated in supplemental figures SF1 and SF2.

### Correlation analysis identifies links between ECM remodeling and tissue biomechanics

In the tumor positive group, Sirus red area in the tumor correlated positively with tumor SWS (*r* = 0.5, *p* < 0.01) and negatively with tumor PR (*r* = –0.6, *p*L< 0.01). Additionally, tumor *φ* correlated strongly with Sirius red area (*r* = 0.6, *p* <L0.001) and moderately with MMP-2–positive area (*r* = 0.5, *p* = 0.01). SWS of the TAL showed moderate positive correlations with Alcian blue area (*r* = 0.4, *p* = 0.04) and MMP-2 area (*r*L=L0.4, *p* = 0.04). Furthermore, SWS of the tumor-bearing livers correlated moderately with Alcian blue area (r = 0.4, *p* = 0.03).

## Discussion

We demonstrated by in vivo mp-MRI and MRE that after inoculation of cancer cells in mouse livers, the entire liver undergoes a biomechanical transition from stiff-viscous to soft-solid behavior within two weeks, preceding any other macroscopic signs of tumor growth.

### Liver biomechanical properties

The observed decrease in liver stiffness during tumor progression can be attributed to the expression of MMP-9, as evidenced by our histological analyses. MMP-9 degrades components of the liver ECM^31^, thereby compromising the overall structural integrity of the liver. The degradation and remodeling effect of MMPs on liver ECM also creates a permissive microenvironment for tumor growth and invasion^32–34^. This is also consistent with the epithelial-mesenchymal (EMT) characteristics of the TIB-75 cells reported in the literature, marked by MMPs upregulation, along with vimentin^35,36^. These changes in ECM structure could be seen by the difference in collagen structure in distinct regions of tumor and liver, in comparison to the control group. The absence of liver fibrosis and inflammatory response to TIB-75 cells suggests MMP-9 expression as a key driver of the observed stiffness reduction^29,37^. The observed reductions in attenuation and fluidity along with restrictions in water diffusivity suggest a link between reduced tissue viscosity and the accumulation of GAGs detected by histology. The hydrophilic nature of sulphated GAGs effectively restricts water mobility^38^ and possibly reduces water diffusion. The influence of GAGs on the in vivo biophysical properties of tumors has previously been reported in the literature^39^, with a reduction of ADC^40,41^ due to water demobilization in gel-like structures. For example, in tumors of central nervous system, GAG accumulation has been discussed as a driver of the transition from stiff-viscous to soft-solid properties^42^. Similar changes in the hepatic microstructure due to the expression of MMPs and GAGs may have caused the observed correlations between stiffness, viscosity and water diffusion shown in Fig.5. Additionally, vessel sprouting—reflected by increased CD31 presence— might also lead to increased liquid fraction and the consequent liver softening. A noteworthy observation is that MMP-9 counts derived from histological analysis exhibited no direct correlation with liver SWS in the HCC group, while the MMP-9 fluorescence intensity was significantly correlated (*r* = –0.6, *p* < 0.01). This finding showed that MMP-9 played a substantial role in shaping the biomechanical environment of the liver. Furthermore, the intensity of MMP-9 expression may serve as a more precise reflection of protein expression levels than the number of counts.

The liver biomechanical properties remained constant in the tumor-negative group, indicating that the observed liver changes were associated with tumor progression rather than injury associated with the surgical procedure of cancer cells implantation. The presence of baseline differences between tumor-positive and tumor-negative animals may reflect recent insights of elevated liver viscoelasticity as a promotor of tumor proliferation^43–46^. Specifically, livers in which tumors proliferated exhibited slightly increased stiffness values compared to those with cicatricial tissue where injected cells did not survive. This suggests that livers with inherently elevated stiffness (high SWS), viscosity (low PR), and fluid-like properties (high φ) provide a tumor-permissive microenvironment that fosters tumor growth^47^.

### Tumor biomechanical properties

Unlike the host liver, the tumor itself exhibited markedly stiff-rigid behavior. Elevated stiffness and reduced water diffusivity of liver tumors likely reflects a higher cellular density and accumulation of collagen in tumor capsule, septae, and peritumoral spaces, as discussed in^15^ and revealed here by histology. Collagen enrichment, especially in HCC, contributes to the formation of a dense fibrotic capsule around the tumor, which acts as a physical barrier that shields cancer cells from immune surveillance promoting invasive growth^48,49^. The dense and compact cellular architecture in tumor masses is not only associated with stiffer properties but also can increase tissue fluidity as seen by the SWS-*φ* correlation analysis in tumors. Increased stiffness was linked to collagen accumulation and matrix maturation, while higher tissue fluidity and reduced viscosity coincided with elevated MMP-2 expression, suggesting that proteolytic activity counterbalances structural rigidity. The inverse relationship between penetration rate and collagen content further highlights the attenuating effect of dense ECM on wave propagation. Regions with high vascular density and mesenchymal transition markers exhibited reduced diffusivity, consistent with increased cellularity and matrix obstruction. In the tumor-adjacent liver (TAL), the association between lower stiffness and MMP-9 expression supports the role of peritumoral proteolysis in regional softening, while the positive correlation of stiffness with glycosaminoglycan content suggests that ECM charge density and hydration may transiently preserve mechanical integrity. Surprisingly, at the same time, attenuation was reduced in stiffer tumors (see SWS-PR correlation in Fig.4), which demonstrates the complementarity of the two viscosity measures, attenuation and fluidity. Attenuation is a loss-related parameter that increases from zero in purely elastic solids to infinity in viscous fluids (vice versa for PR), while fluidity as defined by equation (1) has two limits, zero reflecting nondispersive elastic properties and rr/2 representing the singularity of maximum dispersive shear waves in a fluid^13^. The observation that tumor fluidity positively correlates with stiffness despite reduced attenuation may be explained by the collective behavior of densely packed but unjammed cell clusters that transmit shear waves while establishing multicellular fluid streams^50,51^. It is known that such collective properties are associated with tumor aggressiveness making MRE attenuation and fluidity potentially valuable prognostic markers for the emergent behavior and motility of tumor cells in vivo^52,53^.

### Tumor-liver biomechanical adaptations

The observed correlations of biophysical parameters between tumor and host liver suggest that a mechanically adapted tumor niche is formed after inoculation with cancer cells (Fig. 6). The potential impact of tumor growth on proximal and distal regions was addressed by investigating the TAL study.

**Fig. 6:**
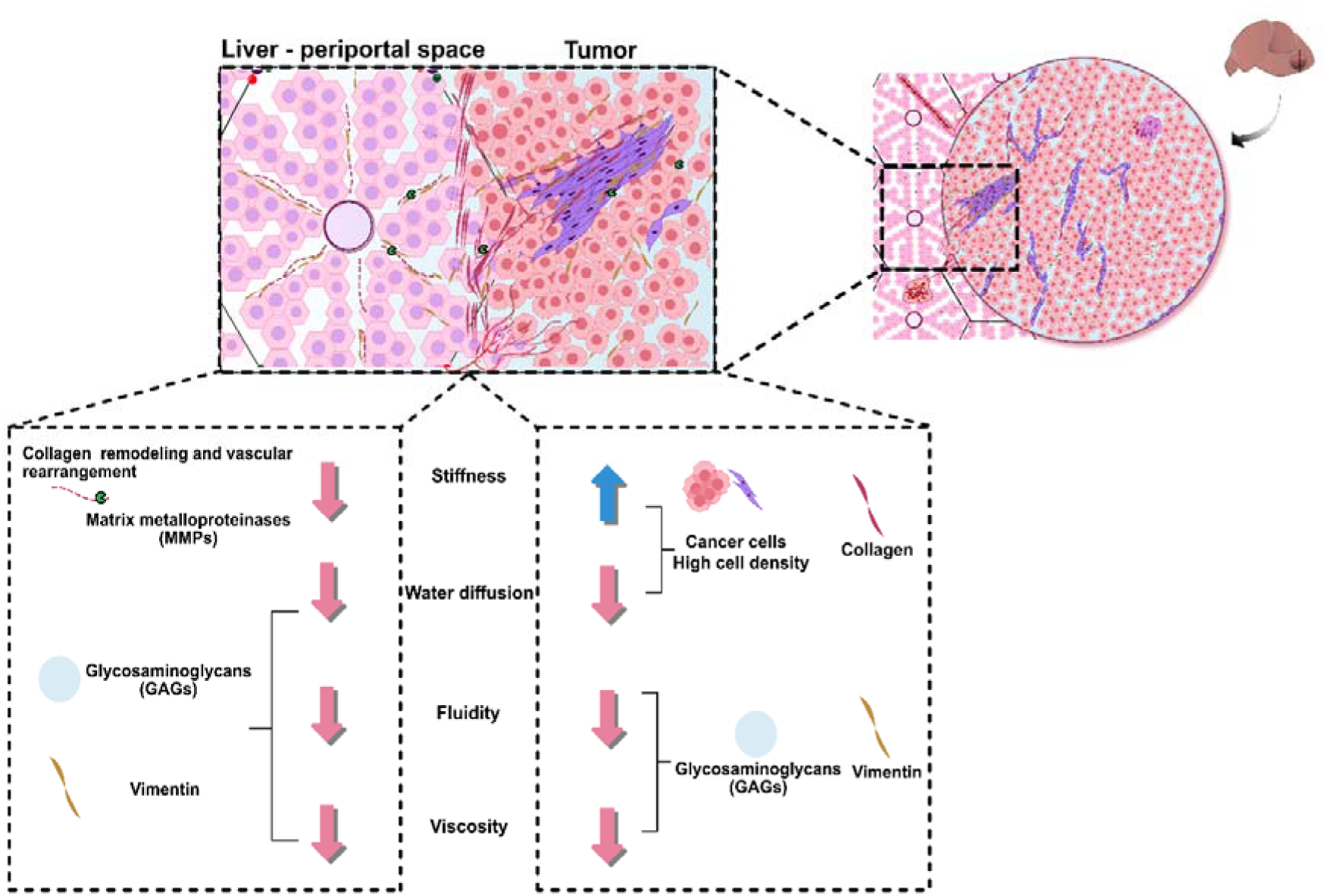
Illustrated summary of liver-tumor microenvironment interactions and biomechanical changes studied herein. A representation of a liver tumor and its surrounding microenvironment. The image illustrates hepatocytes and cancer cells. Extracellular matrix (ECM) is remodeled through metalloproteinases causing liver softening. Furthermore, by forming large polar networks that interact with water molecules, glycosaminoglycans (GAGs) expression and vimentin infiltration are associated with reductions in viscosity and water diffusion. On the other side, the tumor is depicted as a dense cluster of cells with GAGs and vimentin accumulation, contributing to increased stiffness and reductions in viscosity and ADC in the tumorous tissue.

In this peritumoral region, histology showed extended collagen deposition originating from the tumor capsule^54^ along with higher GAG and vimentin content compared to the rest of the liver. Consistent with this result, a transition of MRE parameters from tumor-specific values to liver values was observed with the TAL zone. Despite the differences between tumor and liver MRE values, the extent of their correlations was an unexpected finding. This observed widespread biomechanical adaptation indicates distant biophysical interactions leading to the expression of ECM components such as vimentin, GAGs, collagen, and MMPs, or the propagation of cellular forces into distant ECM^55^. Because vimentin is compartmentalized in both extra– and intracellular spaces, it strongly modulates the coarse-grained biomechanical properties in cell-rich tissues such as liver and HCC^56,57^. As recently reported, vimentin provides structural support that facilitates tumor cell migration and metastasis^58,59^. Mixed with actin gel, vimentin reduces the loss angle of the shear modulus, prompting it as a key regulator of tissue fluidity adaptations between the tumor and liver^60^. Regardless of the mechanism behind the observed biomechanical adaptation, the early signature of tumor cell activity in the liver well before macroscopic tumor growth opens many doors for prognostic medical imaging and novel therapeutic concepts in HCC.

This study has limitations. First, the mouse model used in the current study established HCC in a healthy liver background and short follow up, whereas the majority of human HCC grow in cirrhotic livers after years of chronic liver disease^61^. The knowledge obtained in a healthy liver background (which applies to approximately 10% human cases) can be extended and compared to that obtained in a fibrotic to cirrhotic host liver, which is planned as a follow-up study in our institution. Second, the histological analysis was only available at the final time point. It would be beneficial to harvest and analyze liver samples before the tumor emerges macroscopically to analyze early structural changes of the liver. Third, a contrast-enhanced MRI sequence was not included in the imaging protocol, which will be implemented in future studies to assess tumor vascular physiology and disease extension.

In conclusion, our study used mp-MRI, MRE and histology in a mouse model to characterize the changes in biophysical properties of the in vivo liver during HCC development and progression. Strikingly, we found a widespread softening and reduction in liver viscosity distant to the site of tumor cell injection and well before any macroscopic lesion became visible in conventional MRI. Furthermore, tumor and liver biophysical parameters were closely correlated suggesting an adaptation of tumor and host tissue properties during HCC progression. These findings can have significant implications for the advancement of clinical imaging markers in HCC that target distant pathomechanisms at earlier stages than assessed by state-of-the-art diagnostic modalities when therapeutic interventions are more effective.

## Methods

### Mouse model of HCC

All experimental protocols were approved by the Animal Care and Use Committee in Berlin (LaGeSo, G0048/19) and performed at Charité – Universitätsmedizin Berlin. Animal testing and research conformed to the guidelines set forth in the EU Directive 2010/63 and complied with the Federal German animal protection laws. Briefly, an immunocompetent orthotopic murine model of HCC was established by a single injection 5L×L10L BNL 1ME A.7R.1 cells (TIB-75 cells, ATCC), suspended in 30 µl of Matrigel (Corning Life Sciences), subcapsularly into the left lateral liver lobe of BALB/c mice. Following established protocols, the injection was performed via laparotomy under isoflurane (1.0 %-2.0%) anesthesia and carprofen analgesia (5 mg/kg)^62^.

### Animal use and experiment design

A total of thirty-six 6-week-old BALB/c female mice (Charles River Laboratories, Wilmington, MA, USA) were used in the study and maintained at the institutional Animal Facility with normal water and food supply ad libitum. To monitor cancer growth and quantify the biophysical properties of the tumor as well as the host liver, in vivo longitudinal MRI was performed one week before tumor inoculation (baseline), then at 2w, 4w and 5/6w post tumor inoculation (Fig.7).

**Fig. 7:**
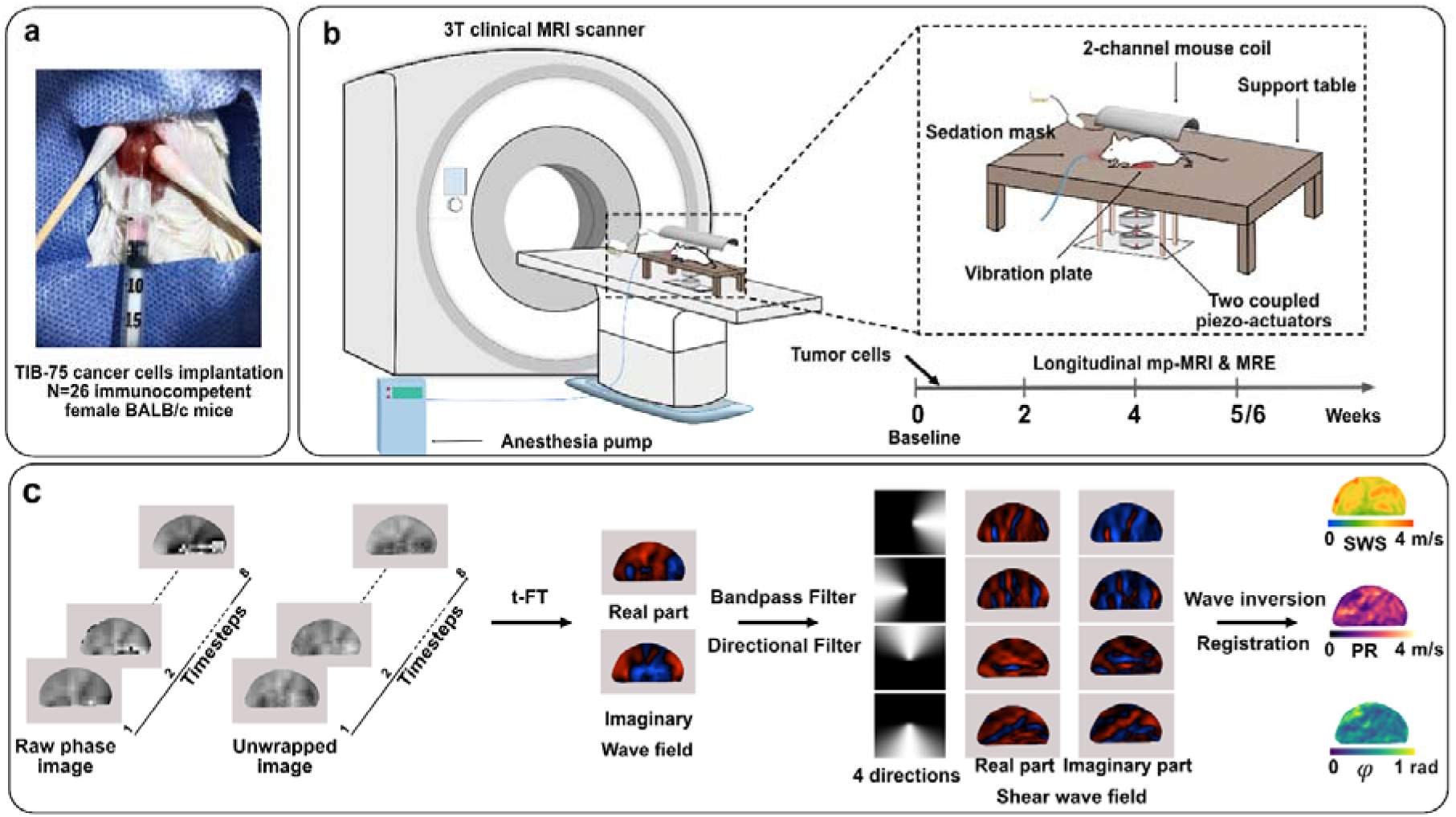
Study design and setup for longitudinal imaging of tumor and adjacent liver tissue in a murine HCC model using MRI and MR elastography. **a** Surgical procedure for cancer cell implantation: laparotomy with microinjection of cancer cells into the left liver lobe. **b** Illustration of the 3 Tesla MR scanner setup, featuring a 2-channel mouse coil and a vibration source. Sedation was maintained via anesthesia pump. The setup included a custom-made sedation mask, precise positioning of the mouse within the 2-channel coil, and a support table. Longitudinal mp-MRI, including T2w, MRE, and DWI sequences, was conducted at various timepoints for all the 36 mice: baseline (one week before tumor inoculation), and two (2w), four (4w), five to six weeks (5/6w) post-tumor inoculation. **c** Vibrations were applied to the mouse abdomen using a vibration plate coupled with two piezo actuators. Data were acquired and phase images were involved in pre– and post-processing steps using the K-MDEV inversion algorithm, enabling the extraction of elastography and diffusion parameters for further biomechanical analysis.

### Mp-MRI and MRE image acquisition

MRI was performed on a 3T MRI system (Magnetom Lumina, Siemens, Germany) equipped with a dual-channel surface coil specifically designed for small animals (mouse heart surface coil array, RAPID Biomedical, Rimpar, Germany). Anesthesia was maintained via inhalation of vaporized 1-2% isoflurane delivered with oxygen through a face mask to ensure the animals remained still during scanning. Animal temperature was controlled with a warm pad during the measurement. The multiparametric MRI protocol included T2w and DWI

#### T2-weighted imaging (T2w)

Anatomical reference images were obtained using turbo-spin-echo (TSE) T2w sequences, acquiring 21 axial slices with an in-plane resolution of 0.25 × 0.25 mm² and a slice thickness of 1.2 mm. Imaging parameters were: echo times (TE) of 77 ms, repetition time (TR) of 2500 ms, and a field of view (FoV) of 150 × 150 mm², with no interslice gap. The total scan duration was 6 minutes and 7 seconds, with no signal averaging (single acquisition).

#### Diffusion-weighted imaging (DWI)

DWI was performed using a multi-shot segmented echo-planar readout (RESOLVE) sequence. A total of 10 axial slices were quired with b-values of 0, 50, 400, and 800 s/mm². The protocol used two separate TEs: 45 ms for data acquisition and 57 ms for navigation in k-space. The TR was set to 2090 ms. The images were acquired with a voxel size of 0.8 × 0.8 mm² in-plane and a slice thickness of 2.0 mm, covering a FoV of 33 × 96 mm². Parallel imaging was performed using GRAPPA with an acceleration factor of 2. ADC maps were automatically processed and generated on the MRI control computer and the values acquired posteriorly analyzed. The scan duration was 6.5 min.

#### Multifrequency MRE

MRE mechanical vibrations were consecutively generated at 300, 400, and 500 Hz frequencies using two synchronized piezoelectric actuators (APA 200, Cedrat Technologies, France). The vibrations were transmitted to the liver through a carbon fiber rod connected to a circular vibration pad (2.5 cm diameter) placed beneath the abdomen of the mouse. Shear wave propagation was imaged using a single-shot spin-echo echo-planar imaging (SE-EPI) sequence with three motion-encoding directions and eight dynamics for each vibration frequency. The frequency of the motion-encoding gradient was 299.40 Hz with an amplitude of 34 mT/m. A total of 21 axial slices (1.2 mm thick) with an in-plane resolution of 1.0 × 1.0 mm² were acquired. The total imaging time of 9.8 minutes included all motion-encoding directions, dynamics, and vibration frequencies. Further imaging parameters were TE = 44 ms, T*r=*2500 ms, and a FoV of 96 × 30 mm².

### Image post-processing

MRE data were processed using the multifrequency wavenumber inversion method k-MDEV^63^. Briefly, a second-order low-pass Butterworth filter (threshold of 2000 mL¹) was applied to the unwrapped phase data for denoising. After Fourier transformation, the wave field data was further processed with a third-order high-pass Butterworth filter (threshold of 650 mL¹) to suppress compression waves. The final inversion step provided maps of SWS (m/s) and PR (m/s), surrogating tissue stiffness and inverse attenuation, respectively. To report the loss angle of the complex shear modulus *φ*, also referred to as fluidity in the literature^10,42^, maps of SWS and PR were converted to fluidity by:

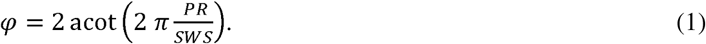

In this study, we report quantitative values of tissue viscoelasticity based on SWS, PR and *φ*, while for the discussion of viscoelasticity changes, colloquial terms of *stiffness, (inverse) attenuation* and *fluidity*, respectively, are used.

### Registration and Masks

Images were first cropped to a uniform size of 100 × 130 mm prior to registration. MR magnitude images were registered to T2-weighted images using rigid alignment. The same transformation was applied to the SWS and PR images. For ADC maps, the DWI with the lowest b-value underwent rigid and affine registration to the T2-weighted images, and the resulting transformation was applied to the ADC maps.

Regions of interest (ROIs) were manually defined in 3D across multiple contiguous slices using T2-weighted and MRE magnitude images for anatomical guidance and were outlined with ITK-SNAP (version 4.2.0)^64^. This volumetric segmentation approach ensured anatomical precision and consistency across the full tumor volume and surrounding liver tissue. Care was taken to exclude liver edges and large vessels. The additional region of interest representing the tumor adjacent liver (TAL) was automatically created by expanding the tumor ROI by 5 pixels on the registered MRE magnitude images. In the tumor-positive group, the liver ROI was defined as the entire liver, excluding the TAL area.

### Histopathological Analysis

#### Tissue harvest and processing

After MR imaging at the final time point (5/6w), animals were euthanized through cervical dislocation in deep anesthesia and in accordance with ethical guidelines. Tumors and adjacent tissues were collected and immediately fixed in 4% formalin (J.T. Baker, Avantor, Radnor, PA, USA) to preserve structural integrity. Tissue samples were then cut into 5 mm sections and embedded in paraffin for histological analysis. Furthermore, liver tissues from ten age-matched healthy mice were harvested and processed in an identical manner to establish a baseline for the histopathological reference.

For routine histopathological assessment, 5μm paraffin sections containing both tumor and surrounding liver tissue were stained with Hematoxylin and Eosin (H & E; Carl Roth GmbH + Co. KG, Karlsruhe, Germany) to assess general tissue architecture and cellular morphology. To evaluate collagen organization and distribution, pentachrome staining was performed following the Doello protocol^65^. Additionally, Alcian blue staining (Morphisto GmbH, Frankfurt, Germany) was utilized to detect GAGs and acidic mucopolysaccharides.

#### Immunofluorescence staining

Immunofluorescence staining was conducted on 5 μm paraffin-embedded tissue sections. Primary antibodies were used to target Metalloproteinase-2 (MMP-2, ab86607, Abcam, Cambridge, UK) and Metalloproteinase-9 (MMP-9, ab283575, Abcam). Vimentin (ab92547, Abcam) was stained to assess mesenchymal markers, while Ki-67 (ab16667, Abcam) and CD-31 (ab182981, Abcam) were used to analyze mitotic proliferation and endothelial cells expression, respectively. Secondary antibodies conjugated to Alexa Fluor 488 and Alexa Fluor 568 (Abcam) were employed for fluorescent imaging.

#### Histological data quantification

Whole-slide images (WSIs) were acquired using high-resolution digitization (ZeBanc, Charité, Germany). Digital annotation masks were manually delineated for each region of interest (ROI) using QuPath v0.4.366 an open-source digital pathology platform.

Hematoxylin and eosin (H&E) sections were analyzed using The StarDist QuPath extension, employed for nuclear detection. Alcian blue and Sirius red were quantified using spectral deconvolution to isolate stain-specific signals, followed by pixel classification (Random Trees) to determine the percentage of stained area and mean optical density. Immunofluorescence markers (MMP-2, MMP-9, Ki-67, Vimentin, CD31) were analyzed based on the corresponding intensity-threshold. Vessel density (vessels/mm²) was assessed based on CD31+ signal.

To further investigate the collagen architecture, Sirius red-stained sections examined with polarized light microscopy with rotatable polarizing filters (0–180° rotation in 10° increments; ZEISS Axio Imager Z1). High-resolution images were acquired under standardized illumination conditions (exposure time = 15 ms, gain = 1×). Birefringence patterning was quantified using the CMM-PR-BRF plugin (ImageJ)^67^, which classifies fibers by maturity based on hue thresholds: mature fibers (red/yellow, 0–40° hue) versus immature fibers (green, 80–120° hue). Fiber spatial organization was assessed using OrientationJ plugin^68^, yielding fiber coherence, a scalar measure of alignment uniformity (0 = isotropic, 1 = fully aligned). The histologic measures were then averaged, with three regions of interest: the tumor, the TAL, and the liver (excluding the TAL). The TAL was delineated as an approximated 1-mm-thick rim encircling the tumor in order to approximate the TAL delineated on the MRI images.

### Statistical Analysis

All statistical analyses and data visualization were performed using R (version 4.4.2; https://www.r-project.org/) and GraphPad Prism 8. Normality of data distribution was assessed using the Shapiro-Wilk test. For normally distributed biomechanical parameters, comparisons across multiple time points and tissue regions were conducted using one-way ANOVA with Tukey’s post-hoc test for pairwise comparisons, while unpaired two-tailed t-tests were used for direct comparisons between tumor-bearing livers and tumor-negative livers. Relationships between normally distributed continuous variables were evaluated using Pearson’s correlation analysis. Alternatively, non-parametric tests were used when analyzing non-normal data. These tests included the Kruskal-Wallis test with Dunn’s post-hoc analysis, the Friedman test, and Spearman’s rank correlation. Group averages were presented as mean ± standard deviation. For histopathological markers, data following Log-normal distribution based on Shapiro-Wilk test were presented as geometric mean (×/÷ SD) for group analysis. A p-value < 0.05 was considered statistically significant.

## Data availability

Requests for access to the protocols and datasets used in this study can be directed to the corresponding author.

## Acknowledgements

This study was funded and supported by SFB1340 Matrix-In-Vision, Heisenberg Grant GU 1726/5-1-Nr: 504873209, FOR5628 and GRK2260 (German Research Foundation, DFG). Authors also acknowledge the Central Biobank Charité (ZeBanC) for their work in digitizing histological slides.

## Author contributions

P.A.D.M., Y.S., and K.K. performed the study design, animal documentation and monitoring, experimental investigations, data collection, statistics, and manuscript preparation. A.H. contributed to project design, animal documentation and monitoring, experimental investigation, and manuscript review. T.H., Y.Z., S.M, R.V.S, J.R and A.A.K. performed histological analysis and manuscript review. L.J.S., J.B., E.E., I.S., and J.G. were involved in grant application, project design and supervision, experimental investigations, animal documentation, data collection, interpretation, and analysis, as well as manuscript writing.

