## Supplemental figures for "Longitudinal in vivo MR elastography reveals whole-liver viscoelastic involvement in a murine model of hepatocellular carcinoma"

**Supplemental Figure SF1**: The following plots illustrate the group statistics of quantitative histological markers in healthy liver, tumor-negative liver, tumor-positive liver, tumor-adjacent liver (TAL), and tumor. Statistically significant post-hoc pairs were indicated with asterisks. **p* < 0.05, ***p* < 0.01, ****p* < 0.001, *****p* < 0.0001.

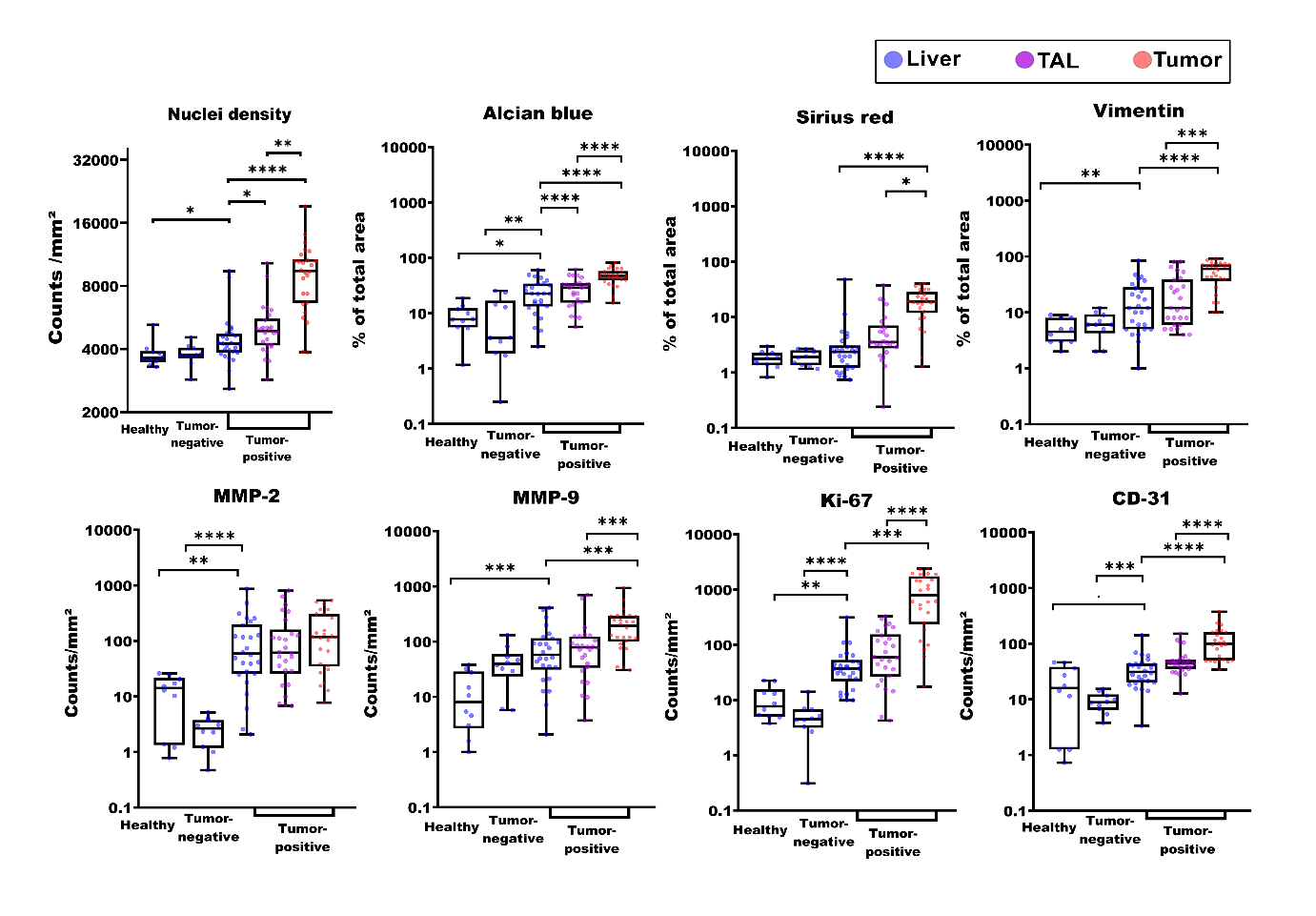

**Supplemental Figure SF2**: The following plots illustrate the group statistics of collagen fiber birefringennce patter and coherency in healthy liver, tumor-negative liver, tumor-positive liver, TAL, and tumor. Birefringence patterning classifies fibers by maturity: mature fibers (red/yellow) versus immature fibers (green). Fiber coherency is a scalar measure of alignment and uniformity (0 = isotropic, 1 = fully aligned). Statistically significant post-hoc pairs were indicated with asterisks.. **p* < 0.05, ***p* < 0.01, ****p* < 0.001, *****p* < 0.0001.

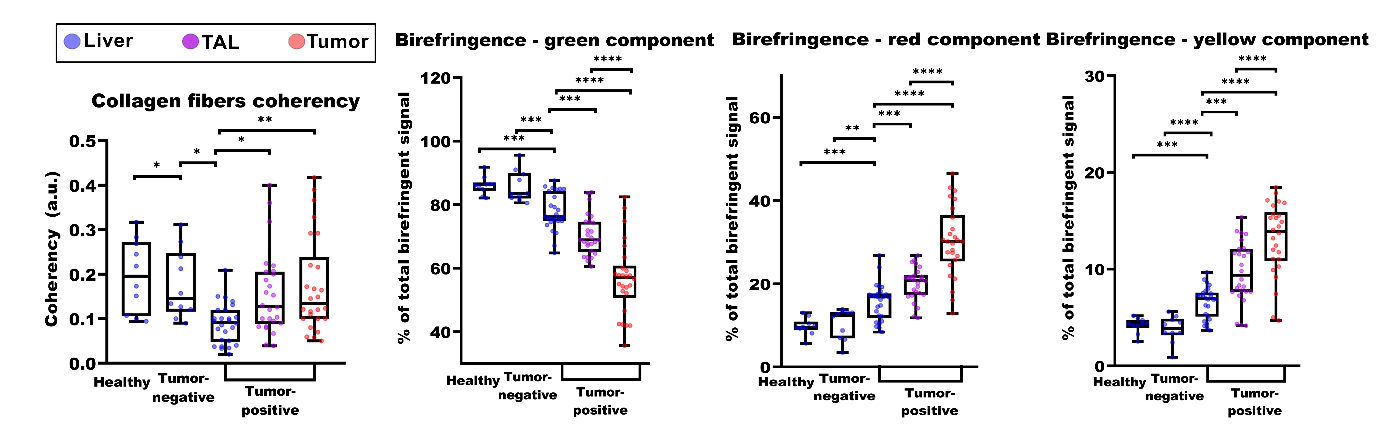
